# Impact of Tissue Sample Preparation Methods on Myelin-Sensitive Quantitative MR Imaging

**DOI:** 10.1101/2025.05.05.652075

**Authors:** Amaya Murguia, Scott D. Swanson, Ulrich Scheven, Andrea Jacobson, Jon-Fredrik Nielsen, Jeffrey A. Fessler, Navid Seraji-Bozorgzad

**Affiliations:** Department of Electrical Engineering and Computer Science, University of Michigan, Ann Arbor, MI, USA; Department of Radiology, Michigan Medicine, University of Michigan, Ann Arbor, MI, USA; Department of Biophysics, University of Michigan, Ann Arbor, MI, USA; Department of Mechanical Engineering, University of Michigan, Ann Arbor, MI, USA; Department of Biomedical Engineering, University of Michigan, Ann Arbor, MI, USA; Department of Neurology, University of Michigan, Ann Arbor, MI, USA

**Keywords:** quantitative MRI, tissue preparation, myelin water imaging (MWI), quantitative magnetization transfer (qMT), bi-exponential T1 mapping, luxol fast blue histology

## Abstract

**Purpose:** Validation of quantitative MRI (qMRI) parameters with histology is often done with ex vivo fixed tissue samples. Freezing is another common form of tissue preservation, but the effects of freezing and thawing tissue on myelin-sensitive quantitative MRI parameters and their correlation with histology require further analysis.

**Methods:** Myelin water imaging, off-resonance RF saturation magnetization transfer (MT), and selective inversion recovery MT MRI experiments were conducted on 14 fresh, thawed, and fixed sheep brain tissue samples to calculate various surrogate measures of myelin content. These measures were compared with luxol fast blue (LFB) histological stain results.

**Results:** Fresh, thawed, and fixed tissue qMRI values correlated well with LFB. Thawed and fixed tissue exhibited modest increases, between 3-32%, for most qMRI parameter values compared to fresh. Histology results showed that thawed samples did not lose tissue integrity from the freezing process.

**Conclusion:** Freezing is a reasonable alternative tissue preservation method to fixation for use in qMRI analysis, but may differentially affect qMRI parameter values in regions with varying myelin content.

## 1 INTRODUCTION

Quantitative MRI (qMRI) has been proposed as a more specific marker of disease pathology than MRI contrast images, and many qMRI methods have been developed to provide quantitative myelin-sensitive metrics ^1^. Many prior qMRI studies have focused specifically on myelinated tissue and myelin-sensitive qMRI, as demyelination is the cause of many diseases, such as multiple sclerosis ^2^, and is involved in many others, such as Alzheimer’s disease ^3^. Quantitative MRI parameters are often validated with histology in fixed tissue samples ^4,5,6,7,8,9^; however, fixation can alter MR tissue properties, including myelin-sensitive properties ^10^, and even immunohistochemistry and histological findings ^11^. Tissue freezing is another method for post-mortem storage that preserves nucleic acid components for genetic analysis on DNA, but a potential concern when using thawed tissue is the effect of the freeze-thaw cycle on cellular integrity, as ice crystals formed during freezing, cold storage, and thawing may destroy tissue components ^12,13^. This study compared the effects of both tissue preparation techniques, freezing and fixation, on myelin-sensitive qMRI parameters using three qMRI methods for analyzing tissue myelin content: myelin water imaging, off-resonance RF saturation magnetization transfer imaging, and selective inversion recovery MT imaging. These methods were chosen since they encompass fundamental contrast mechanisms that are widely accepted as being sensitive to myelin.

Myelin water imaging (MWI) is a common method for analyzing tissue myelin content, and is based on the principle that water molecules within the myelin sheath interact more frequently with molecules of oligodendrocyte membranes, leading to shorter T_2_ times than axonal or extracellular water. The MRI signal is a superposition of water signals in different compartments, leading to multi-exponential decay ^2^. The standard MWI experiment collects multiple spin echoes and fits the curve with non-negative least squares regression to determine the relative signal contribution from each T_2_ component. T_2_ spectra typically have two peaks corresponding to myelin water and free water ^14,15^. The fraction of the signal below a T_2_ cutoff between the two peaks is defined as the myelin water fraction (MWF), which has been histologically validated and accepted as a measure of myelin content ^4,5,6,7,16,17^. However, it has been validated with fixed tissue samples, and fixation has been shown to decrease the T_2_ values of water in the samples ^18,19,20,21,22^ and increase MWF ^10,16,23^. A study using thawed samples found that freezing and thawing tissue decreased water content, but did not significantly affect T_2_ ^24^. Another found that fresh and thawed samples exhibited similar signal behavior when imaged with ultrashort-T_2_ techniques, but the thawed samples had consistently lower signal magnitudes than the fresh samples ^25^. The effects of freezing and thawing tissue on MWF have not been widely studied.

Magnetization transfer (MT) is another common technique used in white matter imaging studies ^26,27,28^. MT in myelin is dominated by exchange of water and macromolecular protons. The macromolecular protons have T_2_ times of order 10 µs and decay too fast to be observed using typical MRI sequences. The MR visible free water protons have much longer T_2_ values. Conventional MT imaging applies an MT preparation pulse far off-resonance from the water signal, directly saturating the macromolecular proton longitudinal magnetization. Proton exchange carries the depleted membrane magnetization to the water molecules, indirectly saturating the free (MR-visible) water magnetization. The magnetization transfer ratio (MTR), the difference between the images obtained with and without the preparation pulse, normalized by the image without the preparation pulse ^29^, has been previously validated in tissue samples for its sensitivity to myelin content ^8,9^. Images acquired after applying preparation pulses at multiple RF power levels and off-resonance frequencies are fit to a quantitative MT (qMT) signal model to calculate the fractional size of the bound pool (F) ^30,31^, which is another accepted measure of myelin content ^32,33,34^. Fixation increases the effect of MT ^22,35,36^, including F ^10^, and freezing can increase MTR ^24^.

Quantitative T_1_ mapping has also been used to study myelination ^37,38,39,40^ and perform qMT analysis ^41,42,43,44,45^. Bi-exponential relaxation of longitudinal magnetization is observed after selective inversion of water proton magnetization. The observed recovery is driven by MT. An inversion pulse selectively inverts water magnetization but not macromolecular proton magnetization. After this pulse, magnetization quickly flows into the inverted water pool from the solid pool. ^46,47^. Inversion recovery data is fit to a biexponential model to calculate the long and short T_1_ components and their corresponding amplitudes. ^37,46^ These parameters can be used to calculate the pool size ratio (PSR), defined as the ratio of the sizes of the macromolecular and free water proton pools ^41^, and has been used as a measure of myelin content ^43,45^. Similar to T_2_ relaxation, fixation decreases the T_1_ value of water ^18,19,20,21,22^, and one study found that freezing tissue also decreases T_1_ ^24^.

In this study, we extended preliminary work ^48,49^ where we conducted MWI and qMRI experiments and luxol fast blue (LFB) histological staining on fresh, thawed, and fixed sheep brain tissue samples to analyze the effects of formalin fixation and freezing on qMRI measures of myelin content. We compared these qMRI parameter values across tissue conditions and correlated them with LFB values.

## 2 METHODS

### 2.1 Samples and equipment

Our process and procedures were compliant with and conducted with the approval of our university’s Institutional Animal Care and Use Committee (IACUC). Seven unfixed, unfrozen ex vivo whole sheep brains were obtained from Nebraska Scientific (Omaha, NE, USA). Immediately after death, the tissue was transferred to a 4°C refrigerator. The time between death and brain extraction was seven days, and the whole brains were shipped to us five days after extraction, with transit time of 12 hours on ice. We sectioned the specimens (see Figure S1 for an illustration) and placed them in the refrigerator at 4°C for up to one day. The total time between death and scanning was 12-13 days, during which time the tissue was at 4°C. The whole brain specimens were cut into *∼*13-mm thick coronal sections; sections at the level of caudate nucleus was placed into histology cassettes of size 40 *×* 26 *×* 13 mm for MR scanning (N=13) One specimen was cut in parasagittal section of the same thickness, resulting in 14 total samples. All samples allowed for examination a variety of tissue types, including WM structures such as the corpus callosum, fornix, cortical white matter, and internal capsel, as well as cortical and subcortical gray matter.

Each cassette was submerged in Fluorinert FC-770 (3M, St. Paul, MN, USA) to minimize B_0_ field inhomogeneity. All specimens were imaged fresh with our entire imaging protocol. Following fresh sample data collection, seven cassettes were placed in a -80°C freezer. After about three weeks the frozen samples were thawed for 48 hours at 4°C before their second round of imaging with the same protocol. The other seven cassettes were placed directly in 10% formalin for 36 hours after the fresh scan, washed in normal saline solution for 12 hours, and scanned again with the same protocol. 36 hours fixation time gives enough time for protein cross linking and tissue preservation, without a significant effect on any subsequent immunohistological methods. This time frame was chosen after discussion with neuropathologists at the Unit for Laboratory Animal Medicine (ULAM) Pathology Core, and proved to be sufficient based on subsequent histological analysis. All samples were placed in 10% formalin and stored at room temperature before histological analysis.

For imaging, the samples were placed in a 40 mm Millipede quadrature MRI coil and inserted into a 7.0 Tesla NMR/MRI scanner (Varian/Agilent, Walnut Creek, CA, USA) with 40 mT/m gradients with a 115-mm inner diameter. The studies were conducted below room temperature at *∼*15-20°C to ensure that the fresh and freshly thawed samples did not decay during the scanning process. Two rounds of B_0_ shimming were performed voxel-wise across the sample using a 3D gradient echo shimming routine. Three of the 14 samples were scanned with a 40 mm saddle coil (Morris Inc., ON, CAN) due to equipment availability; statistical analyses in Section 2.5 accounted for this difference.

### 2.2 Myelin water imaging

Multi-echo spin echo (MESE) data were collected using a multi-echo multi-slice (MEMS) 2D slice-selective sequence with the following parameters: TR = 4000 ms, 64 echoes, echo spacing = 5 ms, matrix size = 128 *×* 128, and Carr-Purcell-Meiboom-Gill (CPMG) phase cycling (custom or default Varian sequence) with two signal averages.

The excitation pulse (90° flip angle) was a 5-lobe sinc pulse with a 1000 µs pulse duration and 5944 Hz bandwidth. The refocusing pulse (180° flip angle) was also an five-lobe sinc pulse with a width of 8000 µs and bandwidth of 5877.5 Hz. Data were acquired for five interleaved slices with thickness = 2 mm, slice gap = 0 mm, field of view = 35 *×* 35 mm^2^, in-plane resolution = 273 *×* 273 µm^2^. The total scan time per sample was 17 min.

The MESE data were analyzed voxel-wise with regularized non-negative least squares (NNLS) regression using the extended phase graph (EPG) formalism to estimate a T_2_ spectrum for each voxel ^14^. The fitting process used 50 T_2_ values log spaced from 5-1000 ms and a regularization parameter of 0.001 to jointly estimate a B_1_ field inhomogeneity scaling factor for each voxel using 16 values linearly spaced from 0.7-1.1; this B_1_ map was then fixed and the T_2_ spectrum was estimated with 500 T_2_ values log spaced from 5-1000 ms and the same regularization parameter ^50,51^. The myelin water fraction (MWF) calculation used a cutoff value of 20 ms ^5^. MWF was calculated for each voxel as the sum of the amplitudes of the points in the T_2_ spectrum up to the cutoff divided by the sum of the amplitudes of the entire spectrum. These values were used to generate a MWF map. We conducted the NNLS fitting with the NNLS package (https://github.com/rdeits/NNLS.jl) in the Julia programming language version 1.8.5 (https://julialang.org). Figure 1 illustrates the steps in the MESE data processing pipeline for a representative fresh sample.

**FIGURE 1.**
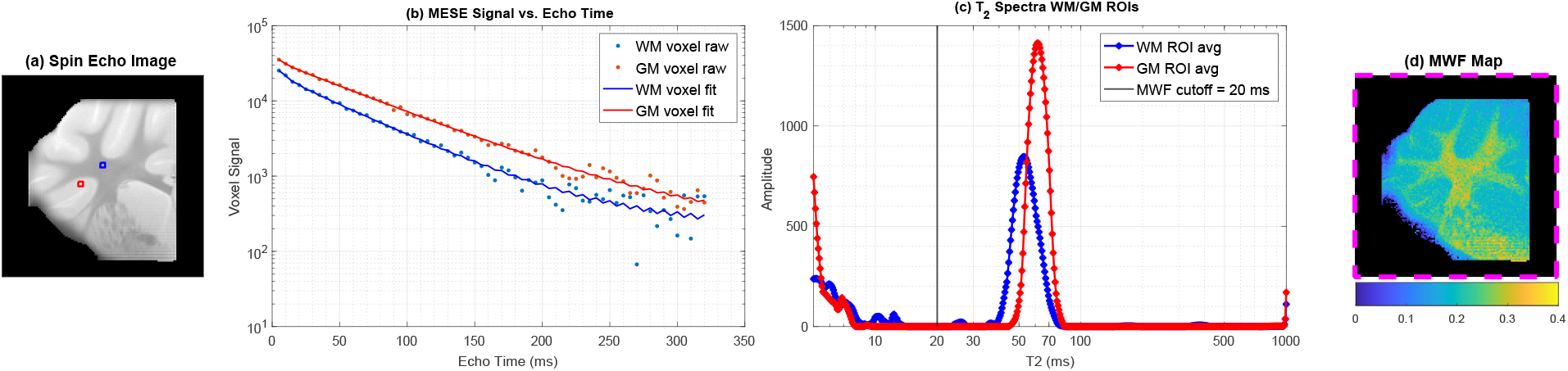
Myelin water imaging analysis for a representative fresh sample ^48^. (a) Spin echo image (TE = 5 ms) marked with WM (blue) and GM ROIs (red). (b) Observed (dotted curve) and fitted (smooth curve) signal decay curves vs. echo time for one WM voxel and one GM voxel in the labeled ROI. (c) Average T_2_ spectra for the WM and GM ROIs. (d) MWF map, calculated using a 20 ms T_2_ cutoff (indicated in (c)). The MWF map is taken as a surrogate measure of myelin content.

### 2.3 Quantitative magnetization transfer

Quantitative MT imaging was conducted using two different sequences and processing methods. The first was steady-state off-resonance RF saturation and MT parameter fitting according to the binary spin-bath model ^30,31^, and the second was transient recovery of selectively inverted water proton magnetization and bi-exponential T_1_ mapping ^41^.

#### 2.3.1 Off-resonance RF saturation

Single-slice MT data were collected via a 2D gradient echo, slice-selective sequence with the following parameters: TR = 120 ms, TE = 3 ms, flip angle = 20°, and matrix size = 128 *×* 128. The excitation pulse (90° flip angle) was a 5-lobe sinc pulse with a 1000 µs width. The MT preparation pulses consisted of a train of 20 Gaussian pulses of duration 1 ms, bandwidth 796 Hz, and duty cycle *∼* 16.7% at 25 off-resonance frequencies from -60kHz to +60kHz at 5 kHz increments and four RF B_1_ RMS amplitudes at about 17.6, 8.8, 4.4 and 2.2 µT with flip angles of 8960°, 4480°, 2240°, and 1120°, respectively. The slice thickness = 2mm, field of view = 35 *×* 35 mm^2^, and in-plane resolution = 273 *×* 273 µm^2^. The total scan time per sample was 26 min.

The magnetization transfer ratio (MTR) was calculated as 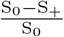, where S_+_ is an image obtained with an MT preparation pulse at 10 kHz and S_0_ is the image obtained at 60 kHz off-resonance. The reported MTR is the average of the MTR maps at +10kHz and -10kHz off-resonance and B_1_ field of 8.8 µT.

A nonlinear parametric voxel-wise fit using a binary spin-bath MT model was performed on the MT data ^30,31,52^. The two-pool model consists of a free water pool, M_f_, with a Lorentzian lineshape and a macromolecular proton pool, M_m_, with a super-Lorentzian lineshape ^53,54,55^. For each voxel, three parameters were fixed. The long T_2_ value of the free water pool, T_2,f_, was assigned a value from a weighted average of of the MWI T_2_ spectrum data for all T_2_ values beyond the 20 ms cut-off. The slow rate of recovery of longitudinal relaxation for the free water pool, R_1,f_, was chosen as the best fit from three values: 0.33, 0.40, and 0.50 s^*−*1 46^; 0.33 s^*−*1^ was chosen. Similarly, the fast rate of recovery of longitudinal relaxation for the macromolecular pool, R_1,m_, was chosen as the best fit from 1 and 1.9 s^*−*1 31,46,56^; 1 s^*−*1^ was chosen. The remaining parameters were estimated: the T_2_ value of the macromolecular pool T_2,m_, the cross-relaxation rate of exchange between the pools R, the ratio of macromolecular to water protons F (referred to as qMT F in this text), and the chemical shift of the macromolecular pool, Δ_cs,m_; see Section S2 in the supporting information for further details about the MT signal model. Corrections for B_0_ and B_1_ inhomogeneities were also included in the model fitting using acquired maps. We conducted the nonlinear fit with MATLAB R2023a (MathWorks, Natick, MA, USA). The qMT F and MTR parameters were used as surrogate measures of myelin content. Figure 2 illustrates the steps in the MT data processing pipeline for a representative fresh sample.

**FIGURE 2.**
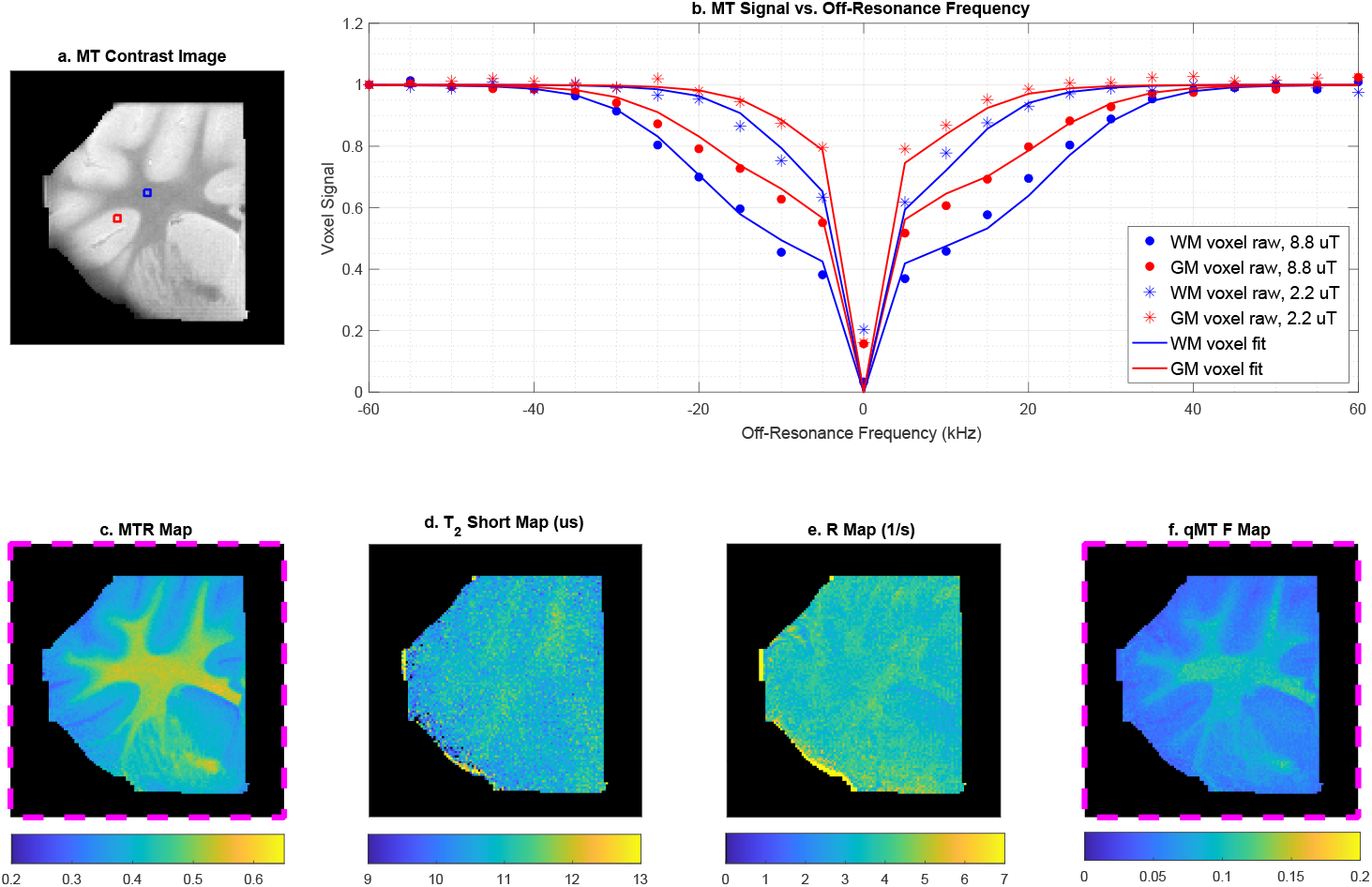
Magnetization transfer analysis for a representative fresh sample ^48^. (a) MT contrast image at 10 kHz off-resonance and second highest power level (8.8 µT) marked with WM (blue) and GM (red) ROIs. (b) Observed and fitted normalized signal curves vs. off-resonance frequencies from -60 kHz to 60 kHz for one WM voxel and one GM voxel in the ROI for two of the four power levels (8.8 and 2.2 µT). (c) MTR map calculated with the average of the -10 kHz and 10 kHz images and 8.8 µT power level. (d) Map of the estimated short T_2_ value corresponding to the macromolecular pool in µs for each voxel. (e) Map of the estimated fundamental rate constant R characterizing the exchange rate between the two pools in seconds for each voxel. (f) Map of the quantitative MT fraction qMT F. Maps outlined in pink (c, f) are taken as surrogate measures of myelin content. The short T_2_ and R maps have low WM/GM contrast compared to prior studies ^31^; this may be due to temperature effects.

#### 2.3.2 Selective inversion recovery

Inversion recovery data were collected using a 2D slice selective inversion recovery (IR) scan with a ramp-sampled spin-echo echo-planar imaging (EPI) readout at 21 inversion times (TI) logarithmically spaced from 10 ms to 5 s with the following parameters: TR = 8000 ms, TE = 36 ms, and matrix size = 64 *×* 64. The excitation pulse (90° flip angle) was a 5-lobe sinc pulse with a 2000 µs width. The refocusing pulse (180° flip angle) was also a 5-lobe sinc pulse with a 1600 µs width. The inversion pulse (180° flip angle) was a hyperbolic secant (HS) adiabatic full passage (AFP) pulse ^57^ with a 4000 µs width and a power of 51 µT. A B_0_ map, which was needed for EPI distortion correction, was acquired separately using a 2D gradient echo slice-selective sequence with multiple TE values at 4, 6, 8, and 10 ms, TR = 100 ms, flip angle = 20°, and matrix size = 128 *×* 128. Data were acquired for five interleaved slices with thickness = 2 mm, slice gap = 0 mm, field of view = 35 *×* 35 mm^2^, and in-plane resolution = 547 *×* 547 µm^2^. The total scan time per sample was 6 min for the EPI IR scan and 1 min for the B_0_ mapping scan.

The IR EPI distorted images were unwarped, upsampled to a 128 *×* 128 matrix size, and registered to the MESE data ^58^. Then a bi-exponential two-pool model fit was performed on the IR data using nonlinear least squares (NLLS) ^41^. Five parameters were estimated voxel-wise: the short and long T_1_ values corresponding to the fast and slow recovery rates, their corresponding amplitudes *a*_fast_ and *a*_slow_, and a B_1_ field inhomogeneity scaling factor. A pool size ratio (PSR) map was calculated as 2*×* the fast relaxing component amplitude *a*_fast_ ^41^ assuming full saturation of the macromolecular pool by the hyperbolic secant pulse; see Sections S3-S4 and Figure S3 for further details. The PSR value was used as a surrogate measure of myelin content. We conducted the nonlinear fit with MATLAB R2023a (MathWorks, Natick, MA, USA). Figure 3 illustrates the steps in the IR data processing pipeline for a representative fresh sample.

**FIGURE 3.**
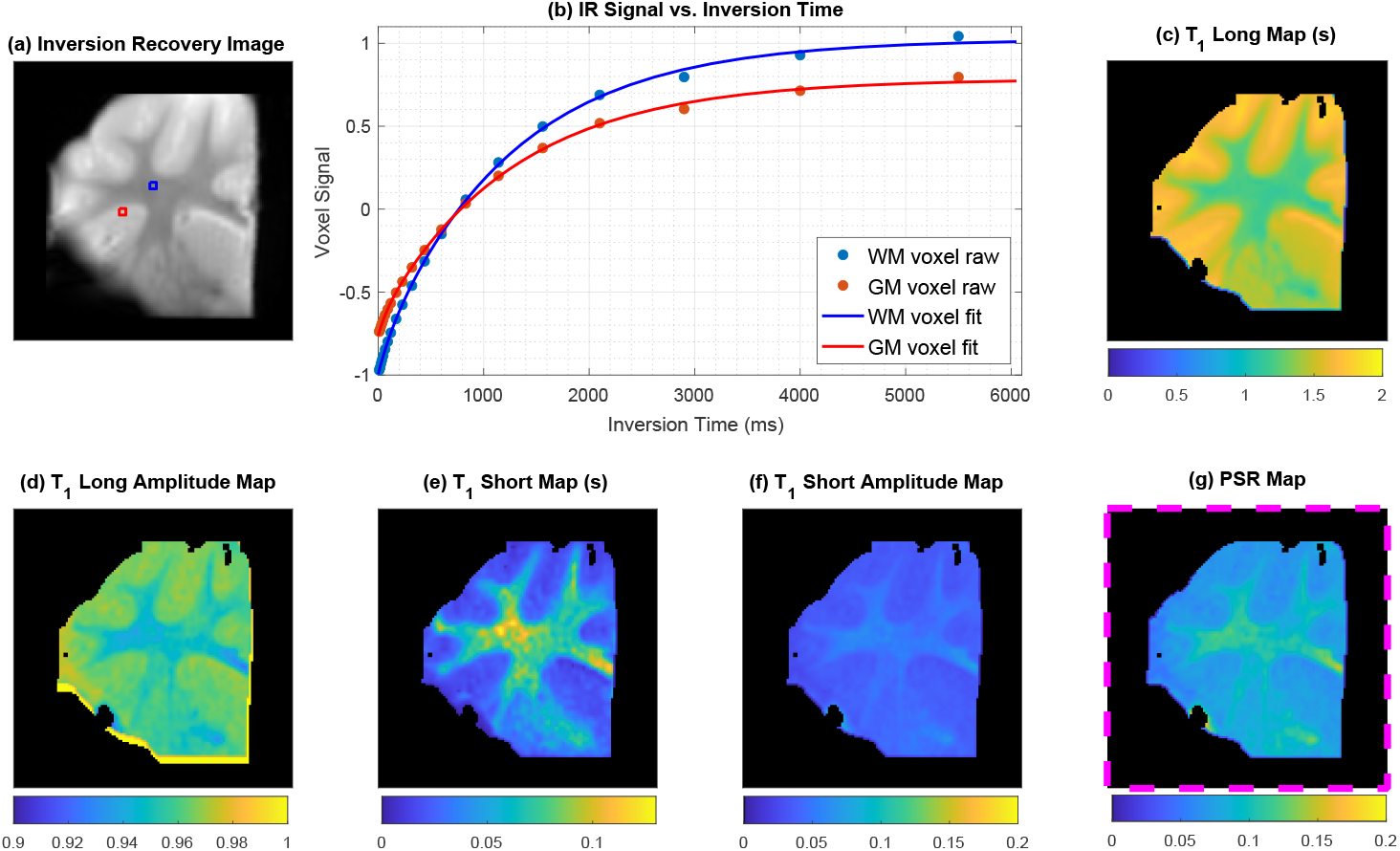
Bi-exponential T_1_ results and analysis for a representative fresh sample ^48^. (a) IR image (TI = 10 ms) marked with WM (blue) and GM (red) ROIs. (b) Observed and fitted signal recovery curves vs. inversion time for one WM voxel and one GM voxel in the ROI. (c) Map of the estimated long T_1_ value for each voxel. (d) Normalized corresponding signal amplitude for the long T_1_ component (*a*_slow_) for each voxel. (e) Map of the estimated short T_1_ value for each voxel. (f) Normalized corresponding signal amplitude for the short T_1_ component (*a*_fast_) for each voxel. (g) Pool size ratio map outlined in pink is taken as a surrogate measure of myelin content.

### 2.4 Histological analysis

After imaging, all samples were immediately placed in 10% formalin to undergo fixation. After fixation, the tissue was bisected along coronal plane with a razor to facilitate paraffin embedding. The top half, based on orientation in the magnet, was stored for further studies, and the other half underwent paraffin embedding, followed by microtomy to obtain a 5 µm slice corresponding to the the center of the original sample. The sample then underwent Luxol fast blue (LFB) staining for myelin visualization and Cresyl Violet (CV) counterstaining for cell nuclei visualization; Figure S2 includes further details. Then they were scanned with a resolution of 0.5 µm/pixel using a Vectra Polaris brightfield whole-slide scanner (Akoya Biosciences, Marlborough, MA, USA). We used the blue channel minus the red channel as a measure of LFB-CV optical density (referred to as LFB in the following text).

### 2.5 ROI and statistical analysis

We used Freeview (https://github.com/freesurfer/freesurfer/tree/fs-7.2/freeview) to label the histology and two corresponding MR center slice images for both tissue conditions (fresh/thawed and fresh/fixed). The images were labeled with WM and GM regions of interest (ROIs), which was done by manually looking for prominent anatomical WM/GM features in both the MR contrast images and histology images. The ROIs were selected using a 4 *×* 4 region in the MR images, corresponding to an area of about 1 mm^2^. The histology ROIs were selected from an 8 *×* 8 region on a downsampled image, corresponding to approximately a 0.25-1 mm^2^ range of tissue. Figure 4 shows a representative sample with corresponding MR and histology WM and GM ROIs labeled.

**FIGURE 4.**
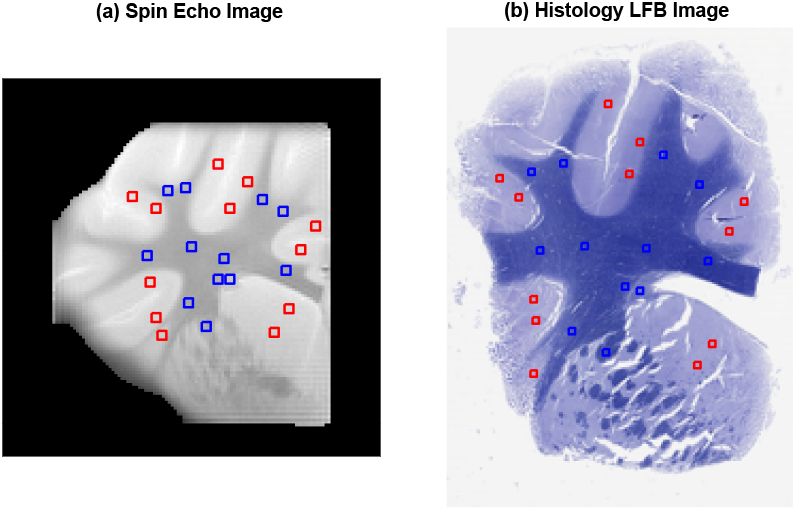
White matter (blue) and gray matter (red) ROIs labeled on a representative fresh sample spin echo image and corresponding histology LFB image.

All statistical analysis was done using R version 4.4.1 (https://www.r-project.org). LFB and qMRI values were averaged across the voxels within each ROI for each sample. These values from all the ROIs were used as individual data points for the subsequent statistical analyses. A mixed-effect linear model with a qMRI parameter as the dependent variable was used; the independent variables include: sample ID modeled as a random effect to account for variability across the samples, tissue condition (fresh/thawed/fixed) modeled as a fixed effect, LFB (or qMRI parameter) modeled as a fixed-effect explanatory variable for regression analysis, and an interaction term between the tissue condition and the explanatory variable when comparing two or more regression terms. A significant interaction term would indicate a difference between the slopes of the regression lines, whereas a significant tissue condition term would indicate a statistically significant intercepts.

The plots were generated as mean-adjusted values generated from the mixed-effect models. The residuals after applying the random effect of sample ID were adjusted to the overall sample mean, which would account for the sample-to-sample variation including the variability in staining and use of different coils. Unpaired t-tests with a Bonferroni corrected p-value threshold of 0.00625 were used for post-hoc pair-wise comparison between the tissue conditions to determine whether the thawed or fixed tissues significantly differed from the fresh samples. Correlation analysis was conducted across ROIs across all samples and also separately for each sample.

Power analysis: for a mixed-effect model, at least 6 samples per tissue preparation method (fresh, thawed, and fixed), and at least three ROIs per region (WM/GM) were needed to achieve a power of 90%, assuming an effect size of at least 10% of the control mean. These calculations were performed using the `simr` package (Version 1.07) function `powerSim` in R. In total, we had 236 ROIs (124 WM, 112 GM) across 14 samples (7 fixed, 7 thawed), with 5 to 24 ROIs per sample.

## 3 RESULTS

Figure 5 shows qMRI parameter maps for two representative samples. Both samples were scanned fresh prior to tissue processing, followed by a second scan after a freeze-thaw cycle for Sample 1 and fixation for Sample 2. We observe similar contrasts in the maps across the different tissue conditions. As expected, a clear contrast was maintained between the WM and GM structures. Visually, freezing or fixation did not significantly affect MR maps or WM/GM contrast. Figure 5 also includes the histology images corresponding to the qMRI maps. The zoomed in images show myelinated axons and intact neurons, oligodendrocytes, and astrocytes for both tissue conditions. However, freezing can cause morphological changes to the tissue, as demonstrated in the Sample 1 LFB image where portions of the sample have cracks. These cracks are also seen in fixed tissue, but based on visual inspection of the histology images, were more prevalent in samples that had been frozen.

**FIGURE 5.**
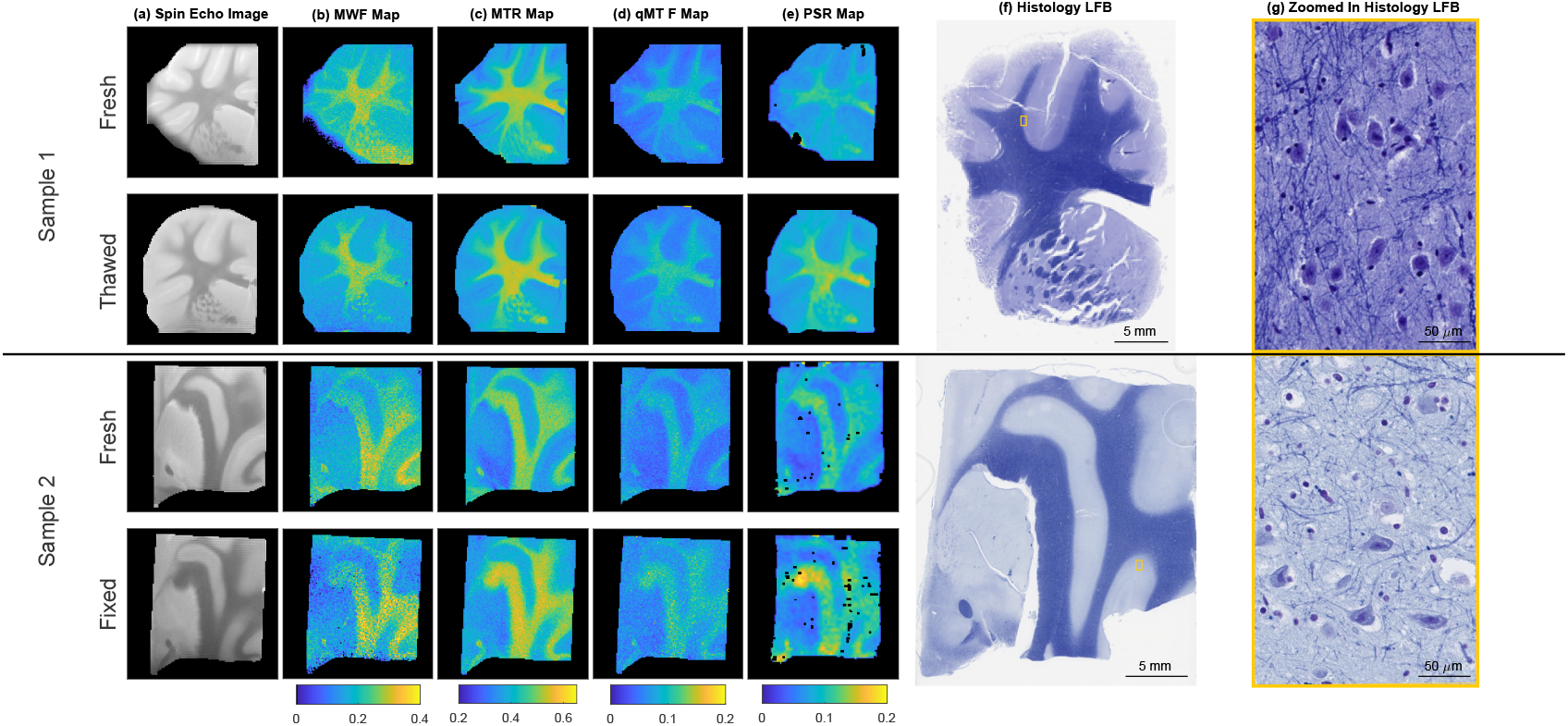
qMRI maps and histology stains for two representative sheep brain tissue samples. Sample 1 (rows 1-2, coronal slice) was scanned fresh and thawed, and Sample 2 (rows 3-4, sagittal slice) was scanned fresh and fixed. (a) Spin echo images at TE = 5 ms. (b) MWF maps calculated with a 20 ms cutoff; this value separated the peaks for most voxels, but black pixel outliers in maps are outlier voxels where the peaks were inconsistent with this cutoff. (c) MTR maps from MT contrast images at average of -10kHz and 10kHz off-resonance and 8.8 µT power level. (d) qMT solid pool fraction maps. (e) PSR maps; black pixel outliers in maps are voxels where the fitting routine produced NaN values. (f) LFB histology for Samples 1-2. (g) Zoomed in views showing cellular integrity.

Figure 6 pools average ROI data from all samples for the four qMRI parameters to show the effect of tissue preparation, separating data based on tissue condition and WM/GM data. The pooled data from the WM ROIs consistently showed a higher value for all qMRI parameters than GM ROIs, supporting the notion that they are sensitive to myelin. Table 1 further quantifies the results in Figure 6 and includes the number of WM/GM ROIs and mean *±* the standard deviation for each qMRI parameter for each tissue condition. The number of ROIs N represents the minimum number of ROIs pooled across samples because some samples are missing data from one or more parameters; see Table S1 for more details. Overall, all qMRI average measures were similar (within *±*0.04) to those of the fresh samples. Tissue preparation seems to differentially affect the qMRI parameters, as some parameters were sensitive to the tissue condition. T-tests detected statistically higher MTR and qMT fraction in WM and GM, and PSR values in WM in both thawed and fixed tissue compared to fresh. The results were comparable across tissue condition for MWF, with the exception that GM MWF values were lower in fixed tissue compared to fresh, a finding that was not observed in the thawed samples.

**TABLE 1.** Quantitative summary of fresh, thawed, and fixed mean qMRI parameter values in Figure 6.

| ROIs | | | Mean ± Standard Deviation<br>P-Values | | | |
| --- | --- | --- | --- | --- | --- | --- |
| Tissue Condition | Type | N | MWF | MTR | qMT F | PSR |
| **Fresh** | WM | 102 | $0.24 \pm 0.035$ | $0.49 \pm 0.026$ | $0.09 \pm 0.011$ | $0.10 \pm 0.012$ |
| | GM | 90 | $0.16 \pm 0.026$ | $0.38 \pm 0.022$ | $0.056 \pm 0.0071$ | $0.070 \pm 0.0096$ |
| **Thawed** | WM | 51 | $0.24 \pm 0.026$<br>$p=0.87$ | **$0.52 \pm 0.027$**<br>**$p=4.1(10^{-10})$** | **$0.11 \pm 0.011$**<br>**$p=1.4(10^{-14})$** | **$0.12 \pm 0.016$**<br>**$p=1.9(10^{-13})$** |
| | GM | 51 | $0.17 \pm 0.020$<br>$p=0.17$ | **$0.39 \pm 0.014$**<br>**$p=0.0011$** | **$0.063 \pm 0.0053$**<br>**$p=8.2(10^{-10})$** | $0.072 \pm 0.0076$<br>$p=0.030$ |
| **Fixed** | WM | 51 | $0.24 \pm 0.054$<br>$p=0.91$ | **$0.53 \pm 0.042$**<br>**$p=4.2(10^{-8})$** | **$0.10 \pm 0.014$**<br>**$p=3.5(10^{-6})$** | **$0.12 \pm 0.018$**<br>**$p=9.5(10^{-8})$** |
| | GM | 39 | **$0.13 \pm 0.028$**<br>**$p=3.2(10^{-9})$** | **$0.39 \pm 0.025$**<br>**$p=0.0026$** | **$0.074 \pm 0.014$**<br>**$p=3.1(10^{-11})$** | $0.072 \pm 0.013$<br>$p=0.16$ |

**FIGURE 6.**
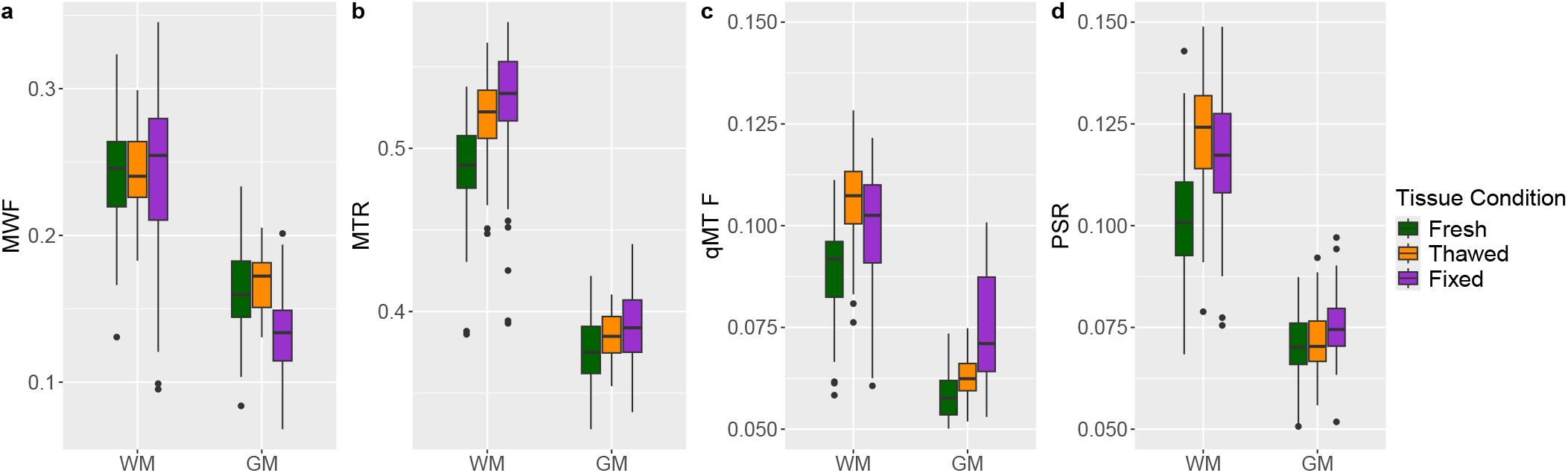
Box plots (interquartile range, whiskers: 1.5-times interquartile range, outliers as dots) of average mean-adjusted MR parameters (across individual samples) from ROIs from all tissue samples. Parameter values are\ consistent with literature values ^5,10,26,45^. For MTR and qMT F, both thawed and fixed mean values were statistically different than fresh mean values for both WM and GM ROIs.

Looking at average WM and GM values of qMRI parameters can mask the effects of tissue processing on estimation of myelin content. Figure 6 exhibits some WM outlier data points with values more in line with GM; for example, some MWF WM points have a very low MWF. We chose ROIs that include a wide range of myelin content, as seen in Figure 4, so the WM-labeled ROIs included some structures with less myelin content, such as the fornix and juxtacortical WM as compared to the corpus callosum. Additionally, tissues with intermediate myelin content in the WM structures may be affected differently by one tissue preparation compared to another. Figure 7 addresses these limitations of the box plots; it shows qMRI values vs. LFB values which demonstrates their correlation and displays intermediate LFB values between the distinct WM and GM clusters. These LFB values include a counterstain; the contribution of the nuclear staining to the overall tissue staining was found to be minimal and relatively uniform given our scale and ROI size as seen in Figure S2 in the supporting information. The graph pools ROI data from samples scanned fresh/thawed and samples scanned fresh/fixed separately; this was done because each pair of scans corresponds to the same LFB data. All qMRI parameters correlated with LFB, and for some plots the qMRI parameters correlated with LFB within the WM ROI subgroups. Table 2 includes quantitative metrics corresponding to Figure 7. It shows the correlation R^2^ values for each qMRI parameter for all ROIs pooled across all samples and for WM and GM ROIs separately, in addition to the range of R^2^ values from correlation analysis conducted on each sample separately The R^2^ values for all ROIs were high (*≥* 0.45) and statistically significant across tissue conditions, as were many of the WM-only and GM-only ROI R^2^ values, especially for MTR; this indicates that LFB accounts for a large variation in the MR signal. The MTR and PSR fresh and thawed lines and MWF fresh and fixed lines had statistically different slopes, and the MWF, MTR, and qMT F fresh and fixed fitted lines had statistically different intercepts.

**TABLE 2.** Quantitative summary of qMRI-LFB correlation in Figure 7.

| ROIs |  |  | MWF | MTR | qMT F | PSR |
| --- | --- | --- | --- | --- | --- | --- |
| Tissue Condition | Type | N | Overall R<sup>2</sup><br>(Range) | Overall R<sup>2</sup><br>(Range) | Overall R<sup>2</sup><br>(Range) | Overall R<sup>2</sup><br>(Range) |
| **Fresh** | WM | 51 | 0.31 | 0.51 | 0.26 | 0.20 |
|  | GM | 51 | - | 0.24 | 0.11 | - |
|  | All | 102 | 0.73<br>(0.75-0.93) | 0.86<br>(0.83-0.98) | 0.79<br>(0.70-0.95) | 0.75<br>(0.63-0.91) |
| **Thawed** | WM | 51 | 0.16 | 0.67 | 0.48 | 0.46 |
|  | GM | 51 | - | - | 0.14 | - |
|  | All | 102 | 0.70<br>(0.64-0.95) | 0.86<br>(0.74-0.98) | 0.84<br>(0.80-0.94) | 0.79<br>(0.70-0.96) |
| Fresh/Thawed Interaction | All | 102 | - | s (p=0.022) | s (p=0.028) | - |
| **Fresh** | WM | 36 | 0.25 | 0.69 | 0.59 | 0.13 |
|  | GM | 24 | - | - | - | 0.44 |
|  | All | 60 | 0.66<br>(0.62-0.88) | 0.94<br>(0.93-0.96) | 0.89<br>(0.86-0.94) | 0.75<br>(0.76-0.78) |
| **Fixed** | WM | 36 | - | 0.15 | 0.11 | - |
|  | GM | 24 | 0.097 | 0.29 | 0.19 | 0.48 |
|  | All | 60 | 0.45<br>(0.42-0.86) | 0.71<br>(0.52-0.98) | 0.64<br>(0.46-0.92) | 0.45<br>(0.32-0.93) |
| Fresh/Fixed Interaction | All | 60 | s (p=0.034)<br>i (p=0.0076) | i (p=0.034) | i (p=8.3(10<sup>-6</sup>)) | - |

**FIGURE 7.**
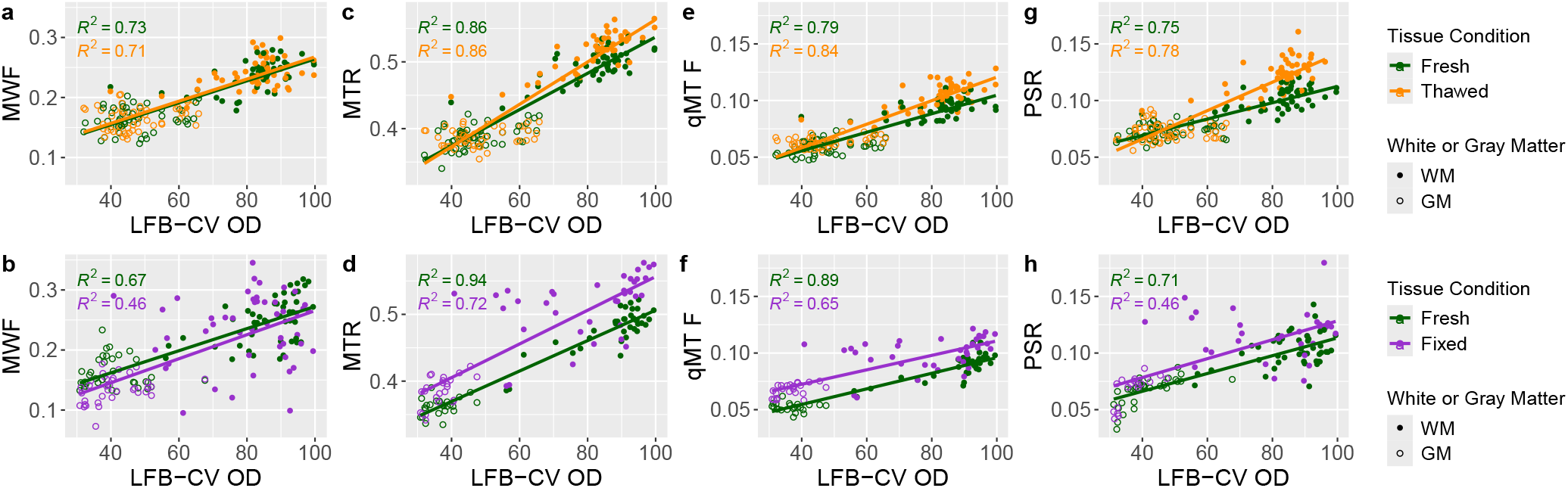
Correlation between qMRI parameter and histology LFB mean-adjusted values across tissue samples for fresh and thawed (Row 1) and fresh and fixed (Row 2) samples. Fresh samples have been separated into two groups based on whether they went on to be frozen or fixed and were not pooled because each pair of scans for one sample corresponds to one LFB image. Fresh, thawed and fixed qMRI parameters showed a strong correlation with LFB OD for all ROIs as indicated by the high R^2^ values.

Figure 8 shows the correlation between each pair of qMRI parameters across ROIs for all tissue conditions. Unlike histology images, where image co-registration is difficult, qMRI-qMRI correlation analysis uses the same ROI coordinates across images. Using similar mixedmodel analysis as with the LFB plots, in subfigures 8c, e, and sub_f_, the slopes and intercepts of the thawed sample data were significantly different from those of the fresh sample data. In subfigures 8d and e the slopes of the fixed sample data were significantly different from the fresh, and in subfigures 8a, d, e, and subf, the intercepts of the fixed sample data were significantly different from the fresh.

**FIGURE 8.**
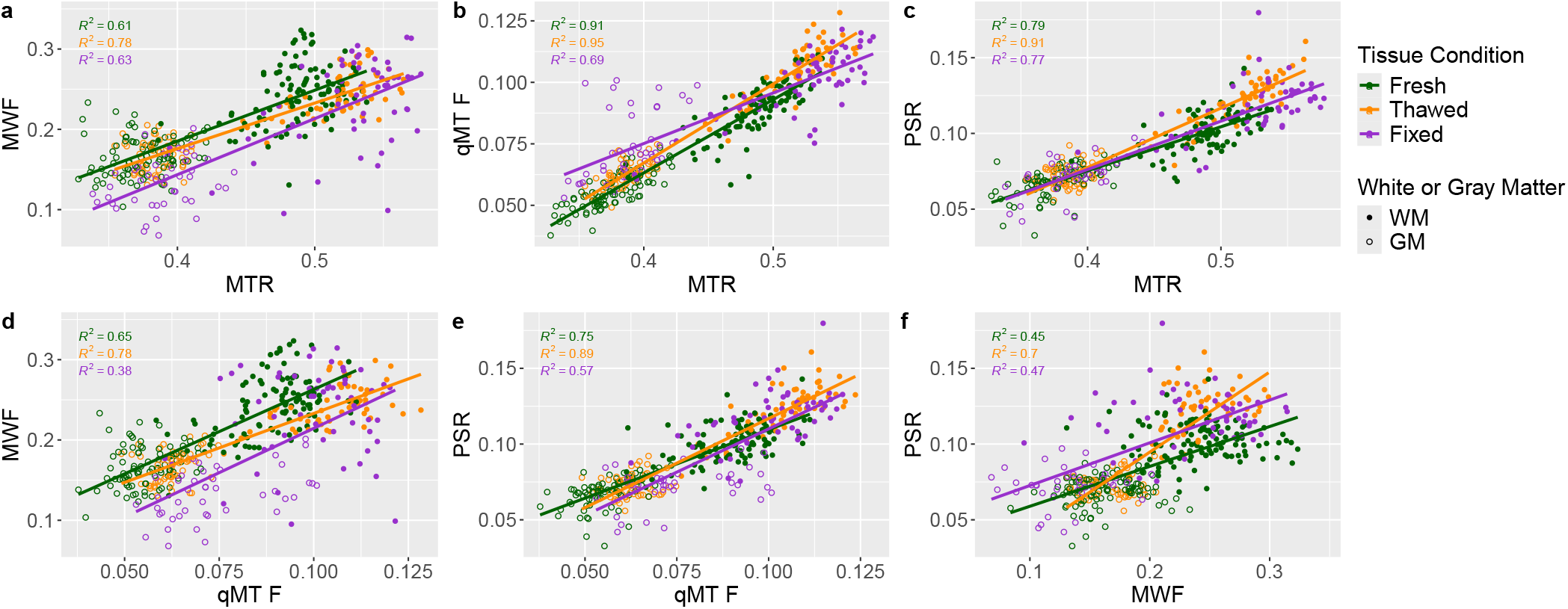
Correlation among each pair of qMRI parameter mean-adjusted values across tissue samples for fresh, thawed, and fixed samples. Fits are similar among tissue conditions for all pairs of parameters, further suggesting that thawed samples are a viable option for qMRI analysis. MTR and qMT F correlate well, likely because both are calculated from the off-resonance RF saturation MT scans. PSR also correlates well with MTR and qMT F, which may result from it also being an MT parameter.

## 4 DISCUSSION

Our findings suggest that alterations in qMRI parameters due to freezing and thawing tissue are comparable to those due to fixation. Although our findings are overall consistent with prior studies, some notable differences are discussed below.

Our ranges of MWF, MTR, qMT F, and PSR values are consistent with those of prior studies ^4,5,6,7,26,44,43,45,59^. We report comparable PSR values in GM, but our PSR values are *∼* 40% higher in WM, which may be due to the water content of tissue, which can decrease ex vivo. It could also be due to temperature effects, as in a separate experiment (not shown) we conducted IR scans of a fixed sample at 19.8°C and 16.3°C and found the PSR value to be *∼* 10 *−* 20% higher at the lower temperature. The qMT-based qMT F and PSR metrics had overall similar means, but the PSR means were *∼* 9 *−* 22% higher, except for the fixed GM case, where the PSR was 3% lower; further comparing these two parameters is an investigation for future studies.

A few groups have studied the effects of tissue fixation on qMRI parameters. In a study of MWF and qMT F values in fresh and fixed 20 mm human spinal cord samples ^10^, Seifert et al. reported an increase of around 40% in MWF for both WM and GM after 24 hours of fixation in 10% formalin compared to the pre-fixation measurement ^10^, which was similar to the observations by Chen et al. ^23^, who studied the effects of fixation on rat cervical spinal cord. Our results, however, show that WM MWF values were not significantly affected by 36 hours of fixation in formalin, and GM values actually decreased by about 19%. Our study design differs from that of Seifert et al. in many aspects; for example, our studies involved different anatomical sites, and we used sheep brain tissue, whereas Seifert et al. used human spinal cord tissue. Our study was conducted at a lower temperature than theirs, which was conducted at room temperature, and MWF has been shown to depend on temperature ^60^. In addition, we submerged formalin-fixed tissue in buffered saline solution for 12 hours prior to scanning, whereas Seifert et al. washed the tissue for multiple 24-hour periods, which could have altered the tissue water content. Since WM and GM may have different susceptibilities to water fluctuations, differences in the duration of the saline wash could potentially affect GM MWF values more than the WM values.

Our MT related measures are consistent with previous studies. We saw a statistically significant increase in MTR and qMT F after fixation, although to a lesser degree than previously reported. For qMT F, Seifert et al. found approximately a 37% increase in WM and a 12% increase in GM after one day of fixation ^10^. We found an approximate 11% increase in WM and a 32% increase in GM after 36 hours of fixation. For MTR, we found a 8% increase in WM and a 3% increase in GM and for PSR, a 20% increase in WM and a 3% increase in GM. The overall smaller change in MT measures due to fixation in our studies could be due to the factors mentioned above for MWF, or the choice of ROIs could contribute to some of the observed differences. We selected ROIs to include less myelinated regions of WM such as the fornix and juxtacortical myelin; this may have affected the mean qMRI parameter values. Our sequences and fitting methods also differed from Seifert et al., which may have affected the results.

A potential explanation for the observed changes in qMT values after tissue fixation is that the fixation expands the spaces between layers of the myelin sheath, which may increase exchange between the macromolecular proton and water pools and increase qMT F ^10^. Formalin cross-links protein lysine residues, which may increase semi-solid content, creating more pathways for MT and increasing MTR.

We also saw significant increases in thawed tissue MTR and qMT F in both WM (6% and 22%) and GM (3% and 13%) and in the PSR in WM (20%) on the same order of the increases from fixation. Evans et al. ^24^ also found that freezing muscle tissue increases MTR, which is consistent with our results, but given the different tissue types, any comparison must be viewed cautiously. Whether the increases in MTR, qMT F, and PSR in frozen tissue are due to similar mechanisms as the effect of fixation requires additional studies with closer inspection of morphological changes with electron microscopy.

Looking at the relationship between qMRI and non-MR measures of myelin content across a wide range of myelin concentration affords additional insight into the effects of tissue preparation on qMRI measures as they relate to other measures such as LFB. Overall, all qMRI parameters correlated well with LFB as in previous studies ^4,5,6,8,9^, even when pooling ROIs from different samples and in WM ROIs alone ^60,61^. As mentioned in Section 3, the slopes of the qMRI vs. LFB plots are generally higher in thawed samples as compared to fresh ones, indicating that the freeze-thaw cycle increases the qMRI signal more for the WM than GM ROIs. This trend reached statistical significance only for MTR and qMT F. In case of the fixed tissue, the intercepts of the fitted lines were affected for MWF, MTR, and qMT F. Tissue fixation appears to affect the WM to the same extent that it affects the GM, thereby increasing the intercepts of qMRI measures of myelin content without as strongly affecting the slopes. We saw a similar trend in the qMRI vs. qMRI plots, which confirmed that the observed trends hold without potential errors due to registration mismatch. The strong correlation in these plots was consistent with prior reports ^10,61^. We also observe a similar trend with MTsat, another qMRI parameter sensitive to myelin content ^62,63,64,65,66,67,68^. Although this parameter has been excluded from the main text, we include MTsat maps and histological correlation analysis in Figures S4-S5.

The correlation analysis results indicate that freezing and fixation may affect qMRI tissue parameters differently; freezing could potentially alter the WM structures that contain thicker and more abundant myelin sheets more robustly than thinner myelinated axons in the GM, whereas fixation could potentially affect the myelination in WM and GM uniformly. We hypothesize that formalin fixation, being a chemical process, may be less dependent on the morphological features of the tissue such as fiber diameter and the number of myelin layers, which may affect WM and GM equally. In contrast, freezing and thawing are thermo-mechanical processes that we conjecture may be affected by the microscopic morphology of the tissue and fiber diameter, thereby having a differential effect on WM and GM. MWF and MTR have been shown to depend on WM fiber orientation and thickness ^61,69^, but further studies are needed to elucidate the mechanisms behind our observations and to test these hypotheses.

Our study unveils many challenges and limitations in evaluating the effects of post-mortem tissue preparation on qMRI measures, which include use of different coils for scanning, lack of strict temperature control, use of semi-quantitative measures of myelin (LFB-CV), variability in the time of death to scan across the samples, lack of stereotactic dissection of tissue and image registration, among others. We attempted to mitigate or account for some of these shortcoming as discussed below. Because we obtained our samples from a vendor, the post-mortem interval ranged from 12 to 13 days; however, care was taken to refrigerate or kept on ice to minimize tissue autolysis, and our histological examination confirmed tissue preservation throughout the process, as seen in Figure 5. Slight difference in postmortem interval, nonetheless, may have affected qMRI results due to increased exchange from breakdown of the cellular membranes.

Additionally, freezing, thawing, and fixation deform the tissue and cause shrinkage and/or expansion, which makes histology-MR image registration challenging, leading us to choose to use manual matching for ROI analysis instead of registration. We selected regions that had clear anatomical demarcation to minimize the effects of slice selections between MR and histology images, but errors in ROI placement undoubtedly remain. These variations may impact findings, such as leading to outliers in the data, impacting why the fixed GM MWF values were lower than their fresh counterparts, and resulting in lower R^2^ values in Figure 7. Figure 8 is immune to these ROI placement errors; however, imperfect unwarping of the EPI data and registration to the MESE data could also have lead to outliers in the PSR values and lower R^2^ values.

Our data indicate that the cells remained intact during the freezing/thawing process of around two weeks, however, the long-term effects of tissue freezing remain unknown. Further experiments such as those in Seifert et al ^10^ are needed to determine how freezing and fixation time affects each of the estimated qMRI parameters. Scanning tissue at lower temperature (*∼*15-20°C) has the advantage of preserving tissue for longer, which is especially important when working with unfixed specimens; however, as mentioned earlier temperature did affect the PSR values, and further experiments are needed to evaluate the effect of temperature on qMRI parameters for different tissue conditions. We did not monitor the temperature as the samples equilibrated from room temperature to the cooler ambient temperature of the magnet at *∼*15°C, so we cannot report the exact temperature, which is a limitation of our study.

Multi-slice MESE MR acquisitions are affected by MT ^51,70^, so in a separate experiment (not shown) we compared qMRI parameters estimated from single and multi-slice MESE and IR acquisitions in a fixed sample. We observed that on average, the MWF/PSR values estimated from the multi-slice acquisitions increased/decreased from the single-slice acquisition by 10 *−* 20%; thus, our use of multi-slice acquisitions is a limitation of our work.

Future studies could also examine the effects of freezing and fixation on MTsat further and on inhomogeneous magnetization transfer (ihMT) imaging, as the ihMT ratio (ihMTR) has been shown to be more specific to myelin than MTR ^60,61,71^. Other modalities such as ultrashort TE (UTE) imaging, quantitative susceptibility mapping (QSM), and quantitative diffusion methods could also be explored.

## 5 CONCLUSION

Thawed and fixed tissue MR parameter values differed modestly compared to fresh values. Both tissue preparations had similar effects on myelin-sensitive qMRI parameter values, and the samples maintained cell integrity, allowing histology to be conducted. Thus, tissue freezing is a reasonable alternative tissue preservation method to fixation for use in qMRI analysis. However, it is important to take into consideration that various tissue preparation techniques may differentially affect regions with varying myelin content or morphology.

## ACKNOWLEDGMENTS

We thank Steven T. Whitaker for sharing his Julia MESE analysis code, and the researchers at the University of British Columbia MRI Research Center for helpful discussions. We also thank the reviewers for comments that greatly improved the paper.

## Data availability statement

Data from all 14 samples will be available here: https://deepblue.lib.umich.edu/data.

## SUPPORTING INFORMATION

Additional Supporting Information may be found online in the Supporting Information section.

### S1 Extra Sample Information

**FIGURE S1** Illustration of the sheep brain dissection and sample selection. The sheep brains were cut into approximately 13 mm coronal sections. The 3rd most rostral section, which was at the level of caudate nucleus, was selected for MR and histological analysis.

**FIGURE S2** LFB + Cresyl Violet plotted against LFB only values (blue minus red channel) across WM ROIs (2 mm diameter circles) show a strong correlation between the two measures, and a slope of nearly 1 from linear regression analysis. As expected, the intercept has a positive value (of 14), indicating a small residual stain for the nuclei in the absence of myelin staining. The plot shows that using a counter-stain has little effect on the estimate of myelin content as compared to LFB alone, at least in ROIs on the order of 1 mm.

**TABLE S1** Quantitative MRI and histology metrics that we had for each sample. Some samples are missing certain scans or did not get sent for histological processing. For example, we do not have MTR or qMT F data for the fresh version of S4, nor do we have histology data for this sample.

### S2 Magnetization Transfer Theory

This section outlines some extra information about the MT signal model that was fit.

### S3 Pool Size Ratio Parameter Calculation

This section outlines some extra information about the calculation of the pool size ratio parameter.

### S4 Comparison of Hyperbolic Secant vs. Gaussian Inversion Pulses

**FIGURE S3** Comparison of hyperbolic secant and Gaussian inversion pulses. (a) IR images (TI = 10 ms) for data collected with the the hyperbolic secant (HS) adiabatic full passage (AFP) inversion pulse ^57^ and Gaussian inversion pulse. (b) Observed and fitted signal recovery curves vs. inversion time for one WM voxel and one GM voxel in the ROI. (c) Map of the estimated long T_1_ value for each voxel. (d) Normalized corresponding signal amplitude for the long T_1_ component (*a*_slow_) for each voxel. (e) Map of the estimated short T_1_ value for each voxel. (f) Normalized corresponding signal amplitude for the short T_1_ component (*a*_fast_) for each voxel. (g) PSR map.

### S5 Additional Myelin-Sensitive Parameter: MTsat

**FIGURE S4** MTR and MTsat maps for two representative samples including the fresh/thawed and fresh/fixed versions of each sample. (a) MTR maps from MT contrast images at average of -10kHz and 10kHz off-resonance and 8.8 µT for fresh/thawed sample. (b) MTR maps for fresh/fixed sample. (c) MTsat maps generated as described above for fresh/thawed sample. (d) MTsat maps for fresh/fixed sample.

**FIGURE S5** Correlation between MTR and MTsat parameters and histology LFB mean-adjusted values across tissue samples for fresh and thawed (Row 1) and fresh and fixed (Row 2) samples. Both MTR and MTsat a show strong correlation with LFB OD for all ROIs as indicated by the high R^2^ values. MTsat exhibits a similar trend to the plots in Figure 7 in the main text based on the mixed-model analysis described in the methods section 2.5. (a-b) exhibit a statistically significant (p*<* 0.05) difference in slopes between the fresh/thawed fits (a: p = 0.022, b: p = 0.0039) and (c-d) exhibit a statistically significant (p *<* 0.05) difference in intercepts between the fresh/fixed fits (c: p = 0.033, d: p= 0.028).

