## Supporting Information for "Impact of Tissue Sample Preparation Methods on Myelin-Sensitive Quantitative MR Imaging"

### S1 | EXTRA SAMPLE INFORMATION

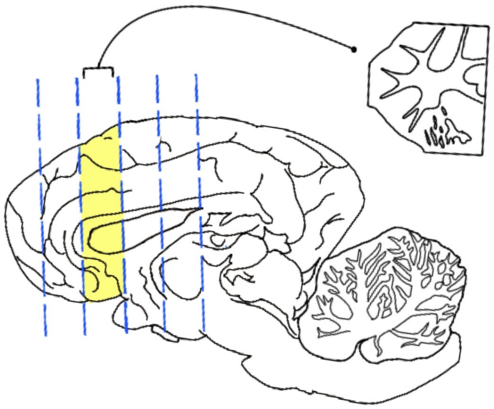

**FIGURE S1** Illustration of the sheep brain dissection and sample selection. The sheep brains were cut into approximately 13 mm coronal sections. The 3rd most rostral section, which was at the level of caudate nucleus, was selected for MR and histological analysis.

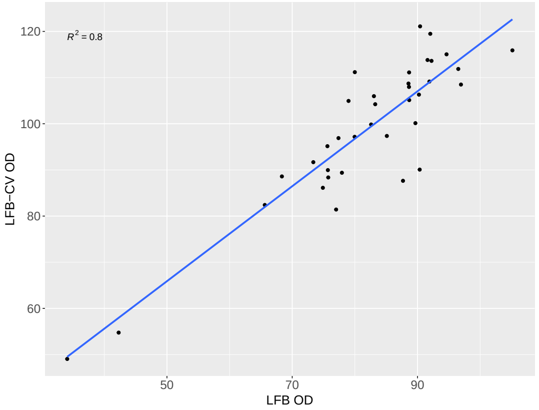


**FIGURE S2** LFB + Cresyl Violet plotted against LFB only values (blue minus red channel) across WM ROIs (2 mm diameter circles) show a strong correlation between the two measures, and a slope of nearly 1 from linear regression analysis. As expected, the intercept has a positive value (of 14), indicating a small residual stain for the nuclei in the absence of myelin staining. The plot shows that using a counter-stain has little effect on the estimate of myelin content as compared to LFB alone, at least in ROIs on the order of 1 mm.

**TABLE S1** Quantitative MRI Parameters and Histology Data Tally for All Samples

| Samples | | Parameter | | | | |
|---|---|---|---|---|---|---|
| ID | Tissue Condition | MWF | MTR | qMT F | PSR | LFB-CV |
| **S1** | Fresh | ✓ | ✓ | ✓ | ✓ | ✓ |
| | Thawed | ✓ | ✓ | ✓ | ✓ | ✓ |
| **S2** | Fresh | ✓ | ✓ | ✓ | ✓ | ✓ |
| | Thawed | ✓ | ✓ | ✓ | ✓ | ✓ |
| **S3** | Fresh | ✓ | ✓ | ✓ | ✓ | ✓ |
| | Thawed | ✓ | ✓ | ✓ | ✓ | ✓ |
| **S4** | Fresh | ✓ | - | - | ✓ | - |
| | Thawed | ✓ | ✓ | ✓ | ✓ | - |
| **S5** | Fresh | ✓ | ✓ | ✓ | ✓ | ✓ |
| | Thawed | ✓ | ✓ | ✓ | ✓ | ✓ |
| **S6** | Fresh | ✓ | ✓ | ✓ | ✓ | ✓ |
| | Thawed | ✓ | ✓ | ✓ | ✓ | ✓ |
| **S7** | Fresh | ✓ | ✓ | ✓ | ✓ | ✓ |
| | Thawed | ✓ | ✓ | ✓ | ✓ | ✓ |
| **S8** | Fresh | ✓ | - | - | ✓ | ✓ |
| | Fixed | ✓ | - | - | - | ✓ |
| **S9** | Fresh | ✓ | ✓ | ✓ | ✓ | - |
| | Fixed | ✓ | ✓ | ✓ | ✓ | - |
| **S10** | Fresh | ✓ | ✓ | ✓ | ✓ | ✓ |
| | Fixed | ✓ | ✓ | ✓ | ✓ | ✓ |
| **S11** | Fresh | ✓ | ✓ | ✓ | ✓ | ✓ |
| | Fixed | ✓ | ✓ | ✓ | ✓ | ✓ |
| **S12** | Fresh | ✓ | ✓ | ✓ | ✓ | - |
| | Fixed | ✓ | ✓ | ✓ | ✓ | - |
| **S13** | Fresh | - | - | - | - | - |
| | Fixed | ✓ | ✓ | ✓ | ✓ | - |
| **S14** | Fresh | ✓ | ✓ | ✓ | ✓ | - |
| | Fixed | ✓ | ✓ | ✓ | ✓ | - |

Quantitative MRI and histology metrics that we had for each sample. Some samples are missing certain scans or did not get sent for histological processing. For example, we do not have MTR or qMT F data for the fresh version of S4, nor do we have histology data for this sample.

### S2 | MAGNETIZATION TRANSFER THEORY

Below outlines some extra information about the magnetization transfer (MT) signal model that was fit [30,31,52]. The water proton magnetization under steady-state RF saturation in a system with a free water proton pool $M_{\mathrm{f}}$ and macromolecular proton pool $M_{\mathrm{m}}$ is given by

$$\frac{M_{\mathrm{sat,f}}}{M_{0,\mathrm{f}}} = \frac{R\left(R_{1,\mathrm{f}} + FR_{1,\mathrm{m}}\right) + R_{1,\mathrm{f}}\left(R_{1,\mathrm{m}} + R_{\mathrm{RF,m}}\right)}{(R_{1,\mathrm{f}} + R_{\mathrm{RF,f}})(R_{1,\mathrm{m}} + R_{\mathrm{RF,m}}) + R\left(R_{1,\mathrm{f}} + R_{\mathrm{RF,f}} + F\left(R_{1,\mathrm{m}} + R_{\mathrm{RF,m}}\right)\right)} \tag{S1}$$

where $\frac{M_{\mathrm{sat,f}}}{M_{0,\mathrm{f}}}$ is the ratio of the free water pool magnetization with and without RF saturation, $R_{1,\mathrm{f}}$ and $R_{1,\mathrm{m}}$ and are the rates of free water and macromolecular proton longitudinal relaxation, $R$ is the cross-relaxation rate, $F$ is the ratio of macromolecular protons to water protons, and $R_{\mathrm{RF,f}}$ and $R_{\mathrm{RF,m}}$ are the rates of RF saturation of the free and macromolecular proton pools given by:

$$R_{\mathrm{RF,p}}(\omega_1, \Delta) = \omega_1^2 g_{\mathrm{p}}(\Delta) \tag{S2}$$

where $\omega_1 = \gamma B_1$ is the RMS amplitude of the applied saturation RF, $\Delta$ is the applied RF frequency in Hz, and $g_{\mathrm{p}}(\Delta)$ is the normalized line shape for pool p (p=f or p=m). The free water pool is given by a Lorentzian function:

$$g_{\mathrm{f}}(2\pi\Delta) = \frac{T_{2,\mathrm{f}}}{1 + (2\pi\Delta T_{2,\mathrm{f}})^2} \tag{S3}$$

and the macromolecular pool is given by a super-Lorentzian function [53,54,55]:

$$g_{\mathrm{m}}(2\pi\Delta) = \pi\sqrt{\frac{2}{\pi}}T_{2,\mathrm{m}} \int_0^1 \frac{1}{\mid 3u^2 - 1 \mid} \exp\left(-2\left(\frac{2\pi(\Delta - \Delta_{\mathrm{cs,m}})\,T_{2,\mathrm{m}}}{3u^2 - 1}\right)^2\right) \mathrm{d}u \tag{S4}$$

where $T_{2,\mathrm{f}}$ and $T_{2,\mathrm{m}}$ are the water proton and macromolecular proton $T_2$ values and $\Delta_{\mathrm{cs,m}}$ is the chemical shift in Hz between the water and macromolecular protons.

The parameters $R$,$T_{2,\mathrm{m}}$, $F$, and $\Delta_{\mathrm{cs,m}}$ were fit on a voxel-by-voxel basis. $R_{1,\mathrm{f}}$ was set to 0.33 $s^{-1}$ [46], $R_{1,\mathrm{m}}$ was set to 1 $s^{-1}$ [31,46,56], and $T_{2,\mathrm{f}}$ was set to a weighted average of the MWI $T_2$ spectrum data for all $T_2$ values beyond the 20 ms cutoff as discussed in the manuscript. $\Delta$ and $\omega_1$ were adjusted by $B_0$ and $B_1$ mapping on a voxel-by-voxel basis.

### S3 | POOL SIZE RATIO PARAMETER CALCULATION

We model the bi-exponential behavior of the inversion recovery $T_1$ mapping data as:

$$\frac{M_\mathrm{f}(t)}{M_{\mathrm{f}\infty}} = a_\mathrm{fast}(1 - 2\exp(-R_1^+ t)) + a_\mathrm{slow}(1 - 2\exp(-R_1^- t)) \tag{S5}$$

where $M_\mathrm{f}(t)$ is the longitudinal magnetization of the mobile protons at time t, $M_{\mathrm{f}\infty}$ is its equilibrium value, $R_1^-$ is the slow recovery rate, $R_1^+$ is the fast recovery rate, and $a_\mathrm{slow}$ and $a_\mathrm{fast}$ are their corresponding amplitudes ($a_\mathrm{slow} + a_\mathrm{fast} = 1$). We calculate a short $T_1$ fraction map (short $T_1$ F) from the fitted amplitudes as: $\frac{a_\mathrm{fast}}{a_\mathrm{fast}+a_\mathrm{slow}} = a_\mathrm{fast}$ [37].

Setting $b_\mathrm{f}^+ = -2\,a_\mathrm{fast}$ and $b_\mathrm{f}^- = -2\,a_\mathrm{slow}$, (S5) can be rewritten in the form of Equation 3 in Gochberg and Gore's paper [41]:

$$\frac{M_\mathrm{f}(t)}{M_{\mathrm{f}\infty}} = b_\mathrm{f}^+ \exp(-R_1^+ t) + b_\mathrm{f}^- \exp(-R_1^- t) + 1 \tag{S6}$$

The pool size ratio can be calculated using the formulation in Gochberg and Gore's paper [41]. We rewrite Equation 6 from Gochberg and Gore's paper using our paper's notation:

$$-2\,a_\mathrm{fast} = \left(\frac{M_\mathrm{f}(0)}{M_{\mathrm{f}\infty}} - \frac{M_\mathrm{m}(0)}{M_{\mathrm{m}\infty}}\right)\left(\frac{p_\mathrm{m}}{p_\mathrm{f}}\right) \tag{S7}$$

where $M_\mathrm{m}(t)$ is the longitudinal magnetization of the macromolecular protons at time t, $M_{\mathrm{m}\infty}$ is its equilibrium value, and $p_\mathrm{m}$ and $p_\mathrm{f}$ are the sizes of the macromolecular and free water proton pools. Solving for the pool size ratio (PSR), which is defined as $\frac{p_\mathrm{m}}{p_\mathrm{f}}$, we get:

$$\mathrm{PSR} = \frac{-2\,a_\mathrm{fast}}{\left(\frac{M_\mathrm{f}(0)}{M_{\mathrm{f}\infty}} - \frac{M_\mathrm{m}(0)}{M_{\mathrm{m}\infty}}\right)} \tag{S8}$$

It follows from (S6) that the ratio $\frac{M_\mathrm{f}(0)}{M_{\mathrm{f}\infty}} = -1$. The hyperbolic secant pulse nearly saturates the macromolecular pool, which makes the ratio $\frac{M_\mathrm{m}(0)}{M_{\mathrm{f}\infty}} \approx 0$. Under these conditions and approximations,

$$\mathrm{PSR}_\mathrm{sech} \approx 2\,a_\mathrm{fast} \tag{S9}$$

Using a Gaussian pulse would cause $\frac{M_m(0)}{M_{\mathrm{f}\infty}}$ to be $\approx 0.88$ according to Gochberg and Gore [41,42]. Under these conditions and approximations,

$$\mathrm{PSR}_\mathrm{gauss} \approx \frac{2\,a_\mathrm{fast}}{1.88} \approx 1.06\,a_\mathrm{fast} \tag{S10}$$

Gochberg and Gore also state that in the case where the macromolecular pool was saturated by the inversion pulse, the PSR values would increase by a factor of $\sim$1.91 [41,42]. Thus,

$$\mathrm{PSR}_\mathrm{sech} \approx 1.91\,\mathrm{PSR}_\mathrm{gauss} \approx 1.91(1.06\,a_\mathrm{fast}) \approx 2.02\,a_\mathrm{fast}\,, \tag{S11}$$

which is consistent with (S9).

### S4 | COMPARISON OF HYPERBOLIC SECANT VS. GAUSSIAN INVERSION PULSES

Our inversion recovery experiments utilized a high power (51 µT) hyperbolic secant pulse[57] that nearly saturated the macromolecular pool, reducing the fast relaxing component amplitude. Based on the calculations in Section S3, in this case the pool size ratio (PSR) would be approximately 2× the fast relaxing component amplitude.

We scanned one thawed sample with two inversion recovery sequences, one with a 51 µT, 4000 µs hyperbolic secant pulse and another with a 7.3 µT, 4000 µs Gaussian pulse and show the results below. The $T_1$ long and $T_1$ short maps do not significantly change depending on the choice of pulse, but their corresponding amplitude maps do. The $T_1$ short amplitude map ($a_{\mathrm{fast}}$) for the Gaussian pulse is about 2× that of the hyperbolic secant pulse, consistent with (S11) in the previous section. Their corresponding PSR maps are consistent with one another and overall with literature values[44,43,45,59], suggesting that the type of pulse used affects the PSR parameter less than it affects the short amplitude.

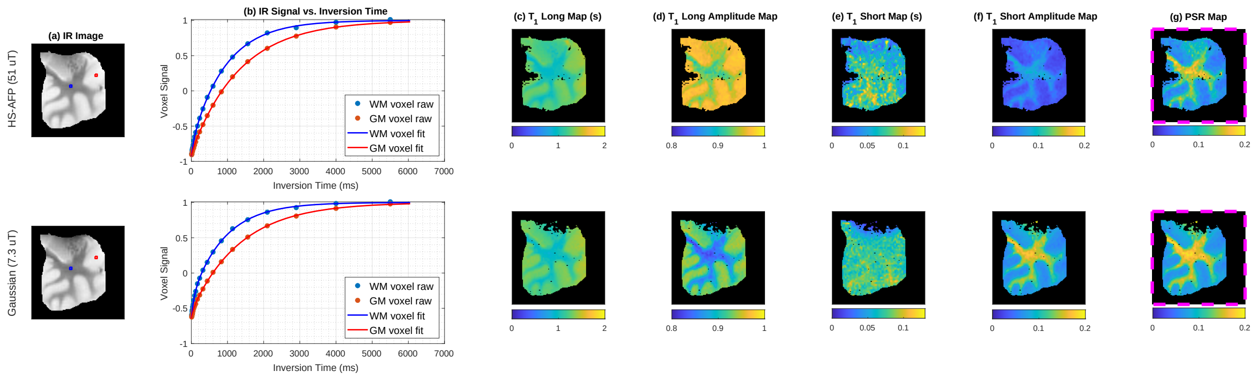


**FIGURE S3** Comparison of hyperbolic secant and Gaussian inversion pulses. (a) IR images (TI = 10 ms) for data collected with the the hyperbolic secant (HS) adiabatic full passage (AFP) inversion pulse[57] and Gaussian inversion pulse. (b) Observed and fitted signal recovery curves vs. inversion time for one WM voxel and one GM voxel in the ROI. (c) Map of the estimated long $T_1$ value for each voxel. (d) Normalized corresponding signal amplitude for the long $T_1$ component ($a_{\mathrm{slow}}$) for each voxel. (e) Map of the estimated short $T_1$ value for each voxel. (f) Normalized corresponding signal amplitude for the short $T_1$ component ($a_{\mathrm{fast}}$) for each voxel. (g) PSR map.

### S5 | ADDITIONAL MYELIN-SENSITIVE PARAMETER: MTSAT

MTsat is another parameter that is sensitive to myelin content with a reduced dependence on $T_1$[62,63,64,65,66,67,68]. These data were added to the supporting information section rather than the main body because to generate these maps we had to generate $T_1$ weighted (T1w) images through simulation since we did not collect this data.

MTsat maps were calculated using Equations 7-8 from Helms et al[62]. The MT weighted (MTw) images acquired at 10 kHz off-resonance and 8.8 µT were used, and the corresponding images acquired at 60 kHz off-resonance were used as the proton density weighted (PDw) images. Apparent $R_1$ ($R_{1\text{app}}$) maps were set to $\frac{1}{T_1\text{Long}}$ estimated in the bi-exponential $T_1$ fitting procedure, and $A_{\text{app}}$ maps were calculated using Equation 7b from Helms et al[62]. Measured $B_1$ maps were incorporated according to Equation 5 in Rahman et al[63]. T1w images of each sample were simulated using TR = 10 ms, flip angle = 50°, our $T_1$ long map, and a proton density map simulated from the PDw image, its TR and flip angle, and our $T_1$ long map. Figure S4 shows MTsat and MTR maps for comparison for a few representative samples. Figure S5 plots MTR and MTsat vs. LFB ROI values to examine their correlation.

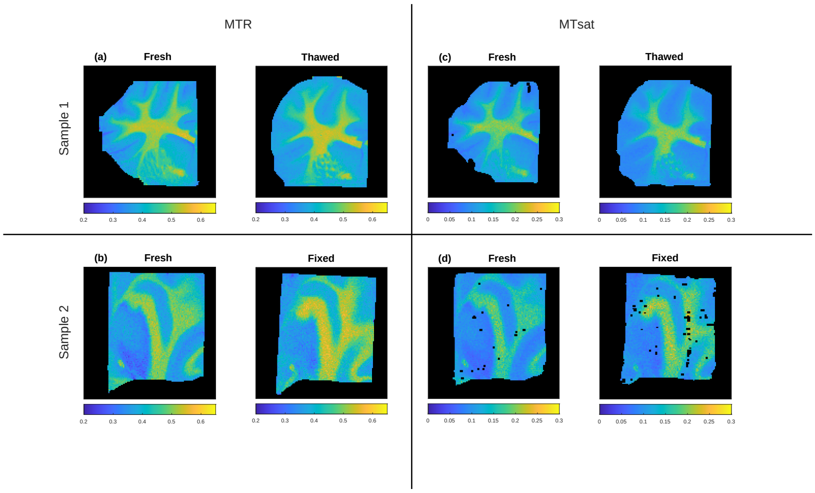


**FIGURE S4** MTR and MTsat maps for two representative samples including the fresh/thawed and fresh/fixed versions of each sample. (a) MTR maps from MT contrast images at average of -10kHz and 10kHz off-resonance and 8.8 µT for fresh/thawed sample. (b) MTR maps for fresh/fixed sample. (c) MTsat maps generated as described above for fresh/thawed sample. (d) MTsat maps for fresh/fixed sample.

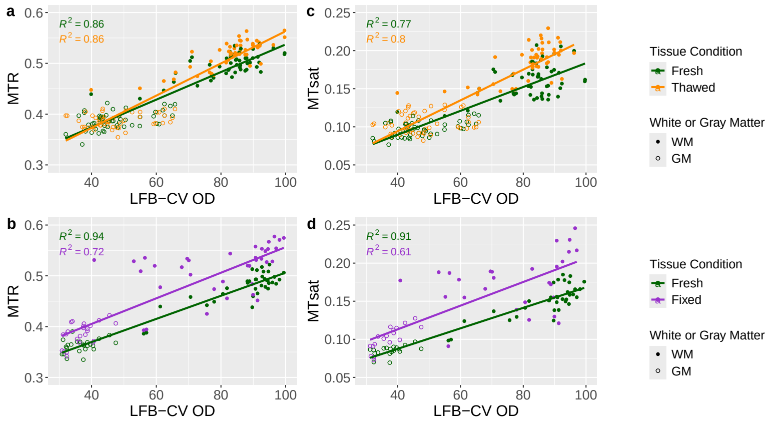


**FIGURE S5** Correlation between MTR and MTsat parameters and histology LFB mean-adjusted values across tissue samples for fresh and thawed (Row 1) and fresh and fixed (Row 2) samples. Both MTR and MTsat a show strong correlation with LFB OD for all ROIs as indicated by the high $R^2$ values. MTsat exhibits a similar trend to the plots in Figure 7 in the main text based on the mixed-model analysis described in the methods section 2.5. (a-b) exhibit a statistically significant ($p < 0.05$) difference in slopes between the fresh/thawed fits (a: $p = 0.022$, b: $p = 0.0039$) and (c-d) exhibit a statistically significant ($p < 0.05$) difference in intercepts between the fresh/fixed fits (c: $p = 0.033$, d: $p = 0.028$).
